# *De novo* transcriptomic characterization of *Betta splendens* for identifying sex-biased genes potentially involved in aggressive behavior modulation and EST-SSR maker development

**DOI:** 10.1101/355354

**Authors:** Wei Yang, Yaorong Wang, Chunhua Zhu, Guangli Li, Hai Huang, Huapu Chen

## Abstract

*Betta splendens* is not only a commercially important labyrinth fish but also a nice research model for understanding the biological underpinnings of aggressive behavior. However, the shortage of basic genetic resource severely inhibits investigations on the molecular mechanism in sexual dimorphism of aggressive behavior typicality, which are essential for further behavior-related studies. There is a lack of knowledge regarding the functional genes involved in aggression expression. The scarce marker resource also impedes research progress of population genetics and genomics. In order to enrich genetic data and sequence resources, transcriptomic analysis was conducted for mature *B. splendens* using a multiple-tissues mixing strategy. A total of 105,505,486 clean reads were obtained and by *de novo* assembly, 69,836 unigenes were generated. Of which, 35,751 unigenes were annotated in at least one of queried databases. The differential expression analysis resulted in 17,683 transcripts differentially expressed between males and females. Plentiful sex-biased genes involved in aggression exhibition were identified via a screening from Gene Ontology terms and Kyoto Encyclopedia of Genes and Genomes pathways, such as *htr*, *drd*, *gabr*, *cyp11a1*, *cyp17a1*, *hsd17b3*, *dax1*, *sf-1*, *hsd17b7*, *gsdf1* and *fem1c*. These putative genes would make good starting points for profound mechanical exploration on aggressive behavioral regulation. Moreover, 12,751 simple sequence repeats were detected from 9,617 unigenes for marker development. Nineteen of the 100 randomly selected primer pairs were demonstrated to be polymorphic. The large amount of transcript sequences will considerably increase available genomic information for gene mining and function analysis, and contribute valuable microsatellite marker resources to in-depth studies on molecular genetics and genomics in the future.

## Introduction

As a freshwater fish species indigenous to Southeast Asia, the Siamese fighting fish *Betta splendens* is one of the most commercially important labyrinth fishes in ornamental fish industry. Among the thousands of aquarium fish species and varieties traded worldwide, *B. splendens* is very popular for its notable value owing to the attractiveness such as variety of color and scale pattern, body shape, fin designs and ease of artificial cultivation in poor quality water [1, 2]. Like some other teleost fishes, *B. splendens* exhibits a distinct reproductive and paternal care behavior [3], whereas there are significant differences in behavior typicality between males and females. Aggressive signaling is an important social behavior of male *B. splendens*, which establish a territory and defend it by displaying higher aggressivity towards intruding conspecifics during reproduction process and post-spawning period [4, 5]. With such specific behavioral typicality, this species has also been gaining increasing popularity as an ideal and useful model organism for behavioral ecology [6], pharmacology [7–10], toxicology [11, 12] and studies examining the biological underpinnings of aggression behavior [3].

Aggressive behavior has received considerable attention and been described in many research projects in vertebrates. Aggression expression is believed to be an intricate and multi-factorially determined biological process. So that, uncovering the basic molecular mechanism of sexual dimorphism of aggression behavior would be very helpful for future behavior-related investigations using *B. splendens* as an experimental model. However, most studies reported on this fish are focused on the behavioral, physiological and metabolic bases of aggression [13], especially including audience effect [14–17], behavioral consistency and variation [18–20], and the effects of sex hormone and steroid hormone mimics [21–23]. To the best of our knowledge, few studies have been conducted on the patterns of gene expression and regulation. The in-depth mechanism underlying the sexual differences of aggressive behavior typicality are still unclear. For this reason, there is an urgent need for more molecular genetics research to illuminate the variance of aggression expression between male and female *B. splendens*.

On the other hand, all wild populations of *B. splendens* are now gravely threatened. With the big progress of commercial breeding and sharp increase of seedling production, the population resource of *B. splendens* has been rapidly declining because of critical hazards such as destruction of natural habitat and contamination from artificially bred fish [1]. It is necessary to pay serious attention to delineating the biodiversity conservation and population structures management in the wild and breeding programs in hatcheries. Therefore, effective and reliable analyzing methods utilizing molecular markers such as simple sequence repeats (SSRs) are strongly recommended.

Unfortunately, the current *B. splendens* genetic information and functional gene sequence resource are extremely scarce. Only 13 genes and 372 DNA and RNA sequences have been deposited in GenBank (analyzed on 26 March 2018). Due to the shortage of basic genomic and genetic resource, the sex-specific genes involved in aggressive behavior are rarely investigated and the molecular mechanism in the sexual dimorphism is poorly understood. Moreover, microsatellite marker development for non-model organisms remains obstructed because of the limitations of time and cost. So far just a few population genetics study has been executed in *B. splendens*, including microsatellite markers development in small quantities [1] and genetic diversity analysis by means of allozyme markers [2]. The scant marker resource severely impedes the research progress of population genetics and genomics in fighting fish. Undoubtedly, both molecular genetics and further investigations on mechanism of aggression exhibition significantly request more genetic background knowledge.

Transcriptome is a small but essential part of the genome and contains a large amount of encoding genes. With improved efficiency, next-generation sequencing (NGS) based transcriptome high-throughput sequencing is widely utilized to obtain large-scale genetic information and generate great amount of transcript sequences and gene expression data rapidly and cost-effectively, especially for non-model species [24, 25]. Currently transcriptome sequencing has been frequently employed to functional gene discovery, gene expression and regulation analysis, screening of large sets of molecular markers [26]. In the present study, Illumina RNA sequencing, Trinity *de novo* assembly and annotation, differential expression analysis, and SSRs screening were firstly performed in *B. splendens* using multiple tissues mixing strategy. This study mainly aimed i) to markedly enrich genetic data and functional gene sequences for gene mining and identification of candidate genes putatively involved in sexual dimorphism of aggression behavior typicality, and ii) to develop polymorphic EST-SSR markers for future research into population genetics and genomics.

## Materials and methods

### Ethics statement

The experimental animals used in this study were artificially cultivated. All animal experiments were approved by the Institutional Animal Care and Use Committee (IACUC) of Guangdong Ocean University, Guangdong, China. All sampling procedures were complied with the guidelines of IACUC on the care and use of animals for scientific purposes.

### Animal materials and samples collection

For transcriptome sequencing, one-year-old mature *B. splendens* (females: n=10; males: n=10) were purchased from a fighting fish hatchery center (Sanya, Hainan, China). All fish samples were acclimated in laboratory for one week before use. Alive fish were sacrificed by decapitation following anesthetization with a tricaine methanesulfonate (MS222) immersion bath. Tissue samples including brain, heart, muscle, liver, kidney, gut, spleen, gill and gonad (testis or ovary) were excised as soon as possible. All samples above were quick-frozen in liquid nitrogen immediately, and then stored at −80°C until RNA extraction. To verify the polymorphism of EST-SSR markers, skeletal muscle tissues were excised from thirty *B. splendens* individuals collected randomly from the broodstock population in the hatchery center. The muscle samples were placed in absolute ethanol and kept at −20°C until genomic DNA isolation.

### RNA extraction, cDNA library construction and sequencing

Total RNA was isolated from each tissue of female and male *B. splendens* using a Trizol reagent kit (Life Technologies, Carlsbad, CA, USA) following the instructions of manufacturer. The extracted RNA was treated with RNase-free DNase I (TaKaRa Biotech Co., Ltd., Dalian, China) to remove residual genomic DNA. The concentration of total RNA samples was quantified by Nanodrop 2000c (Thermo Scientific, Wilmington, DE, USA) using the absorbance value at 260 nm, and the purity was determined by OD_260/280_ (accept range: 1.8-2.0). The 18*S* and 28*S* ribosomal bands stained with ethidium bromide (EB) on a 0.8% agarose gel were utilized for RNA integrity assessment. After pooling the RNA samples from different tissues in equal quantity (approximately 1 μg each tissue), the male and female pooled RNA samples (approximately 5 μg each sex) were delivered to Gene Denovo Biotechnology Co., Ltd. (Guangzhou, Guangdong, China) for cDNA library construction and sequencing. mRNA was purified using an Oligo-dT Beads Kit (Qiagen, Hilden, Germany) and cDNA libraries were constructed following the protocol for Illumina RNA sequencing, respectively. The two cDNA libraries were sequenced on Illumina HiSeq™ 2000 sequencing platform (Illumina, Inc., San Diego, USA) and paired-end (PE) reads with a length of 125 bp were generated.

### *De novo* assembly

By means of SOAPnuke v1.5.0, the raw sequencing data was quality-controlled with the parameters “−l 10 −q 0.5 −n 0.05 −p 1 −i”. The raw reads were filtered to generate high quality data via processes including the removal of adapter sequences, reads with ambiguous sequences (N) more than 10% and sequences with more than 20% low-quality bases (quality value < 20). Subsequently, the clean reads with high quality were *de novo* assembled by the Trinity RNA-Seq Assembler (version: r20140717, http://trinityrnaseq.sourceforge.net/) with default parameters [26]. At first, clean reads were assembled by Inchworm using a greedy k-mer based approach, resulting in a collection of linear contigs. Then, the abundant contigs were built into de Bruijn graphs through k-1 overlaps by Chrysalis. Finally, the fragmented de Bruijn graphs were trimmed, compacted and reconciled to final linear transcripts using Butterfly. The redundant final linear transcripts sequences were eliminated and the longest ones were defined as unigenes [27].

### Annotation and classification

Functional annotation was performed by sequence alignment against public databases using BLAST 2.2.26+ software with an E-value cut-off of 1E-5. All assembled unigenes were submitted to sequence homology searches against protein databases including NCBI non-redundant protein (Nr), EuKaryotic Orthologous Group (KOG), Swiss-Prot, Kyoto Encyclopedia of Genes and Genomes (KEGG) by BLASTx. Sequences with the highest similarity scores from the databases were defined as the functional annotation for the related unigenes. Further analysis on the annotation results were carried out to obtain the Gene Ontology (GO) functional results by Blast2GO software [28]. And then GO term classification and visualization were accomplished by WEGO statistical software [29]. The KEGG pathway annotation was analyzed by the KOBAS v2.0 software for pathway categories [30]. Moreover, assembled unigenes were also aligned to KOG database to predict and classify possible functions.

### Differential expression analysis

The reads per kb per million reads (RPKM) method was utilized to conduct the comparison analysis of the difference in gene expression in this study. With the ability to eliminate the detrimental effects of different gene lengths and sequencing levels on the calculation of gene expression, the RPKM values could be used directly for comparison analysis of the difference in gene expression between variant samples [31]. Differential expression analysis of unigenes between the two sexes was carried out by the edgeR package [32]. *P* value was adjusted by means of false discovery rate (FDR). FDR value <0.05 and |log_2_ (Fold Change)| >1 was set as the threshold for significantly differential expression. The obtained differentially expressed genes were also defined as sex-biased genes (SBGs) in this study.

### Validation of SBGs by qRT-PCR

Twelve SBGs putatively associated with aggression behavioral regulation were selected to validate the results of RNA-seq analysis by qRT-PCR. Based on the transcript sequences derived from the transcriptome, specific primers were designed using Primer 5.0 (S1 Table). Total RNA was extracted using TRIzol Reagent and treated with DNase I to avoid genomic contamination. The first-strand cDNA was synthesized from approximately 1 μg RNA using a RevertAid First Strand cDNA Synthesis kit (Fermentas, Vilnius, Lithuania). qRT-PCR was performed on a LightCycler 480 system (Roche, Basel, Switzerland) using SYBR Premix Ex Taq II (TaKaRa Bio Inc., Shiga, Japan). The PCR amplification conditions were as follows: 95 °C for 1 min followed by 40 cycles of 95 °C for 10 s, and 60 - 64 °C (depending on different genes) for 30 s. Three independent biological triplicates and two technique repeats were performed for each sample. The reference gene β-actin was used to determine the relative expression. The relative gene expression levels were calculated using 2^−ΔΔCt^ method. ANOVA analysis was performed by SPSS 17.0 statistical program (SPSS Inc., Chicago, USA). The difference with *P* < 0.05 was considered significant.

### SSR loci search and primer design

SSR loci in assembled unigene sequences were detected using the perl program MIcroSAtellite (MISA, http://pgrc.ipk-gatersleben.de/misa/) [33]. Due to some difficulties in distinguishing genuine mononucleotide repeats from polyadenylation products and single nucleotide stretch errors generated by sequencing, mononucleotide repeats were excluded in this study. A minimum of six, five, four, four and four contiguous repeat units were applied to identify di-, tri-, tetra-, penta- and hexa-nucleotide SSR motifs, respectively. The primer pairs flanking each SSR locus were designed by Primer 5.0 software [34].

### SSR polymorphism examination

Genomic DNA was extracted from the muscle samples using a marine animal tissue genomic DNA kit (Tiangen Biotech, Beijing, China) following the manufacturer’s protocols. The quality and quantity of extracted DNA was determined by NanoDrop2000 spectrophotometer (Thermo Scientific, Willmington, DE, USA) and 1.0% agarose gel electrophoresis. The DNA samples were diluted to a final concentration of 20 ng/μL with ddH_2_O and stored at −20°C for PCR analysis. PCR was performed on the C1000™ Thermal Cycler (Bio-Rad, CA, USA) in a total volume of 10.0 μL. Each reaction tube contains 1.0 μL of the DNA template (20 ng), 0.2 μL of each primer (10 μmol/L), 5.0 μL of the 2×EasyTaq PCR SuperMix (Invitrogen, California, USA), and 3.6 μL of ddH_2_O. PCR cycling conditions consisted of initial denaturation at 94 °C for 5 min, followed by 30 cycles of 45 s at 94 °C, 40 s at the annealing temperature, and 40 s at 72 °C, and a final extension at 72 °C for 5 min. PCR products were separated on a 8.0% non-denaturing polyacrylamide gel, and allele sizes were estimated according to the pBR322 DNA/MspI marker (Tiangen Biotech, Beijing, China). The number of alleles (*N*_a_), expected heterozygosity (*H*_e_), observed heterozygosity (*H*_o_), and polymorphism information content (*PIC*) per SSR locus were calculated by PowerMarker v3.25 software [35].

## Results

### Sequencing and assembly

The male and female cDNA libraries were sequenced using Illumina sequencing technology. The raw sequencing data were uploaded to the SRA databases of NCBI with accession numbers SRX2558711 for male and SRX2558712 for female. Intotality, 108,416,100 raw PE reads (13.55 Gb sequencing data) were obtained. The quality control of raw reads resulted in 105,505,486 clean PE reads (53,062,092 for female and 52,443,394 for male), which is equal to 13.19 Gb sequencing data. The Q20 percentage and GC content of the clean data were 97.315% and 50.635%, respectively (Table 1). These high-quality clean reads were *de novo* assembled subsequently, and a total of 69,836 unigenes with a mean length of 1093.52 bp and an N50 length of 2040 bp were generated, representing a total of 76.37 Mb sequences (Table 1). For sequence length distribution, all assembled unigenes ranged from 228 bp to 20,412 bp. Exactly 37,510 (53.71%) unigenes were >500 bp in length, 22,876 (32.76%) were >1000 bp and 11,290 (16.17%) unigenes were >2000 bp in length (Fig 1).

**Table 1.** Statistical summary of sequencing and assembled of *B. splendens* transcriptome.

| Sequencing |  |  |  | Assembly |  |
| --- | --- | --- | --- | --- | --- |
| Item | Female library | Male library | Average | Item | Value |
| Raw reads | 54,600,768 | 53,815,332 | 108,416,100 | Unigenes | 69,836 |
| Raw bases (bp) | 6,825,096,000 | 6,726,916,500 | 13,552,012,500 | GC content (%) | 50.66 |
| Clean reads | 53,062,092 | 52,443,394 | 105,505,486 | Total length (bp) | 76,366,868 |
| Clean bases (bp) | 6,632,761,500 | 6,555,424,250 | 13,188,185,750 | N50 length (bp) | 2,040 |
| Q20 percentage (%) | 97.18 | 97.45 | 97.315 | Mean length (bp) | 1,093.52 |
| GC content (%) | 50.89 | 50.38 | 50.635 | — | — |

**Fig 1.**
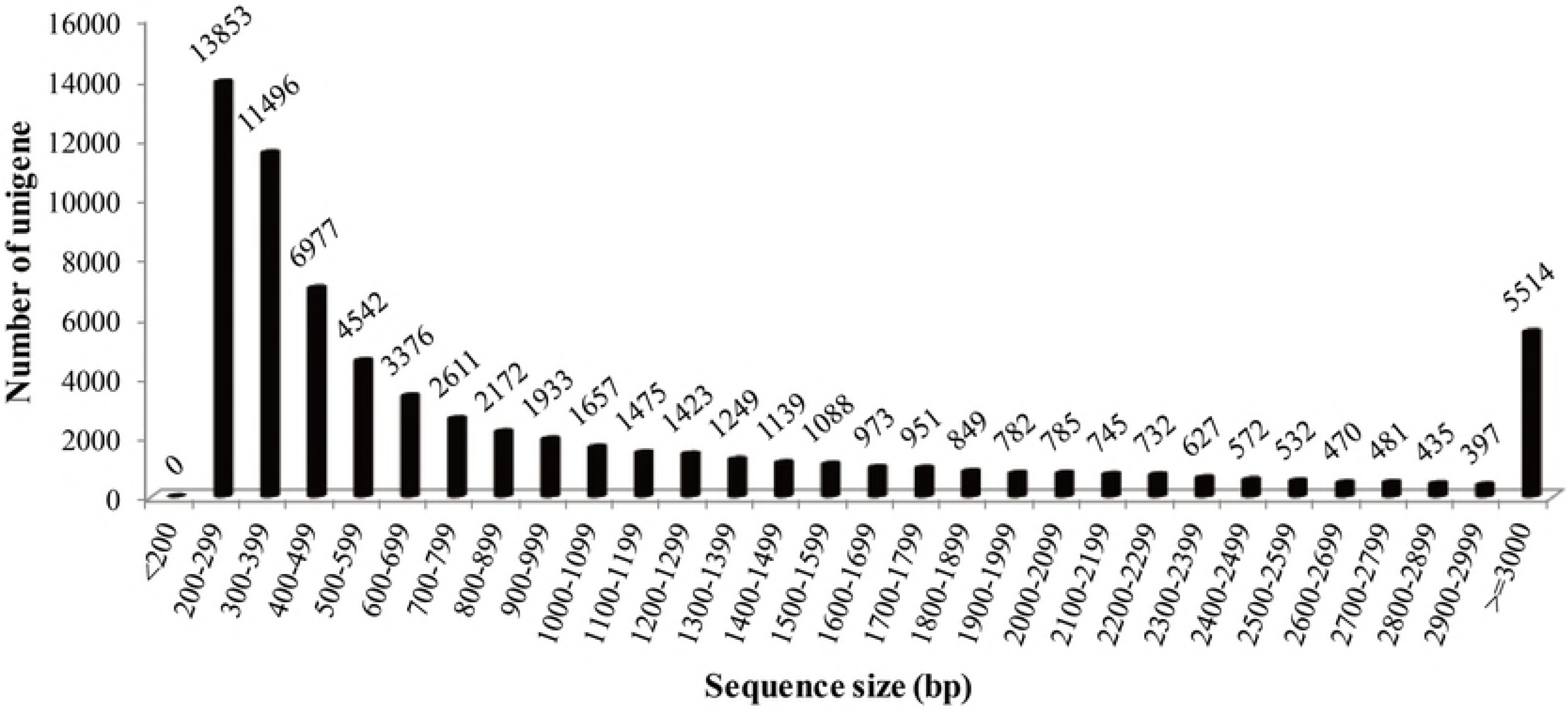
The sequence size distribution of assembled unigenes derived from *B. splendens* transcriptome.

### Functional annotation

Unigene sequences were annotated through BLASTx search against public databases. A total of 35,751 (51.19%) unigenes were annotated in at least one of queried databases. Out of these unigenes, 35,707 (51.13%), 29,791 (42.66%), 23,978 (34.33%) and 16,956 (24.28%) unigenes showed significant similarities to the subject proteins in the Nr, Swiss-Prot, KOG and KEGG databases, respectively. 14,608 (24.28%) unigenes could be annotated in all databases simultaneously (Fig 2A). The E-value distribution of blast analysis showed that 72.36% unigenes exhibited significant homology (below 1E-50) in Nr database and 16.74% homolog sequences ranged from 1E-20 to 1E-50 (Fig 2B). Meanwhile, 79.19% and 33.22% of the sequences exceeded 70% and 90% similarity, respectively (Fig 2C). A further analysis on the matching sequences showed that the homologous genes came from 304 species. Of which, 27.51% of the unigenes had the highest homology to genes from *Larimichthys crocea*, followed by *Stegastes partitus* (23.59%), *Oreochromis niloticus* (8.15%), *Notothenia coriiceps* (4.70%), *Maylandia zebra* (3.89%) and *Neolamprologus brichardi* (3.15%), above six species accounting for over 70% of the annotated unigenes (Fig 2D).

**Fig 2.**
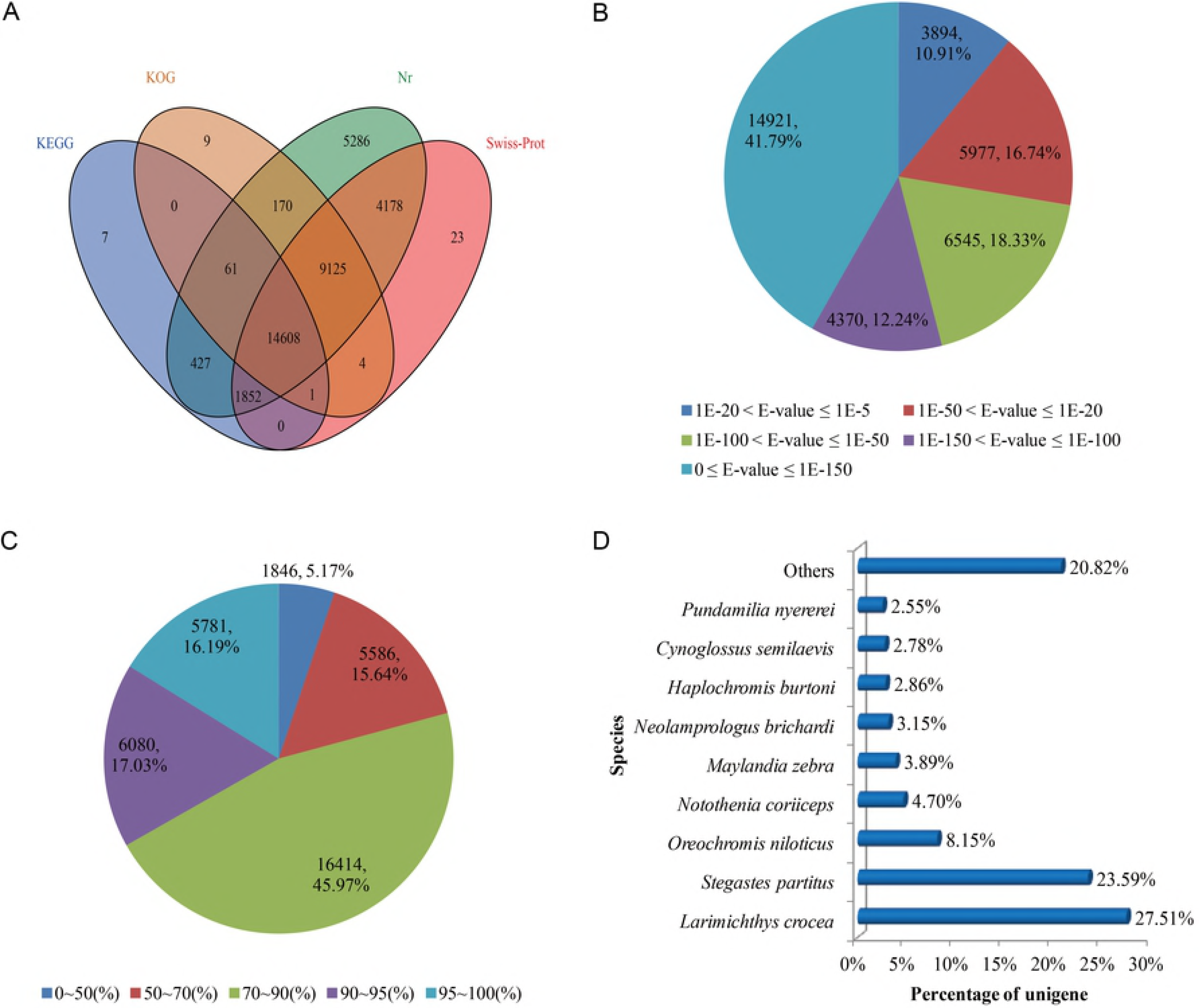
Statistical summary of assembly and annotation of unigenes. (A) Venn diagram of unigenes annotation results against Nr, Swiss-Prot, KOG and KEGG databases. The number in each colored block indicates the number of unigenes annotated by single or multiple databases. (B) E-value distribution of BLAST search against the Nr database for each unique sequence (E-value 1E-5). (C) Identity distribution of BLAST hits for each unique sequence. (D) Species distribution of the BLAST hits for the assembled unigenes (E-value 1E-5). The top nine matching species were shown.

### GO, KEGG pathway and KOG classification

For understanding the function of the unigenes, GO terms were assigned to each sequence using Blast2GO, and then GO classification and visualization were finished by WEGO tools. In totality, 16,954 (24.28%) unigenes annotated in GO database with one or more GO terms were assigned into 56 level-2 functional groups (Fig 3A, S2 Table). Among which, 55,254 (55.43%) genes comprised ‘biological process’ as the largest category, followed by ‘cellular component’ (26,491; 26.58%) and ‘molecular function’ (17,930; 17.99%). The GO terms ‘cellular process’ (9,249; 9.28%) and ‘single-organism process’ (8,137; 8.16%), ‘cell’ (5,393; 5.41%) and ‘cell part’ (4,939; 4.95%), ‘binding’ (8,796; 8.82%) and ‘catalytic activity’ (5,513; 5.53%) were the primary and secondary largest clusters in the three main GO categories, respectively.

The unigenes were subjected to KEGG pathway analysis to further investigate the biological pathways activated in *B. splendens*. A total of 16,956 (24.28%) unigenes were assigned to six main categories, including 240 different KEGG pathways and 35,751 genes (Fig 3B, S3 Table). Among the main categories, the largest one was ‘human diseases’, which contained 9,809 (27.44%) KEGG-annotated genes, followed by ‘organismal systems’ (9,460; 26.46%), ‘metabolism’ (6,188; 17.31%), ‘environmental information processing’ (4,474; 12.51%), ‘cellular processes’ (3,825; 10.70%) and ‘genetic information processing’ (1,995; 5.58%).Moreover, 55.53% of sequences in the top ten hit pathways were involved in ‘global and overview maps’, ‘cancers: overview’ and ‘signal transduction’, whereas the others were related to pathways such as ‘signaling molecules and interaction’, ‘cell motility’, ‘cellular community’ and ‘development’.

**Fig 3.**
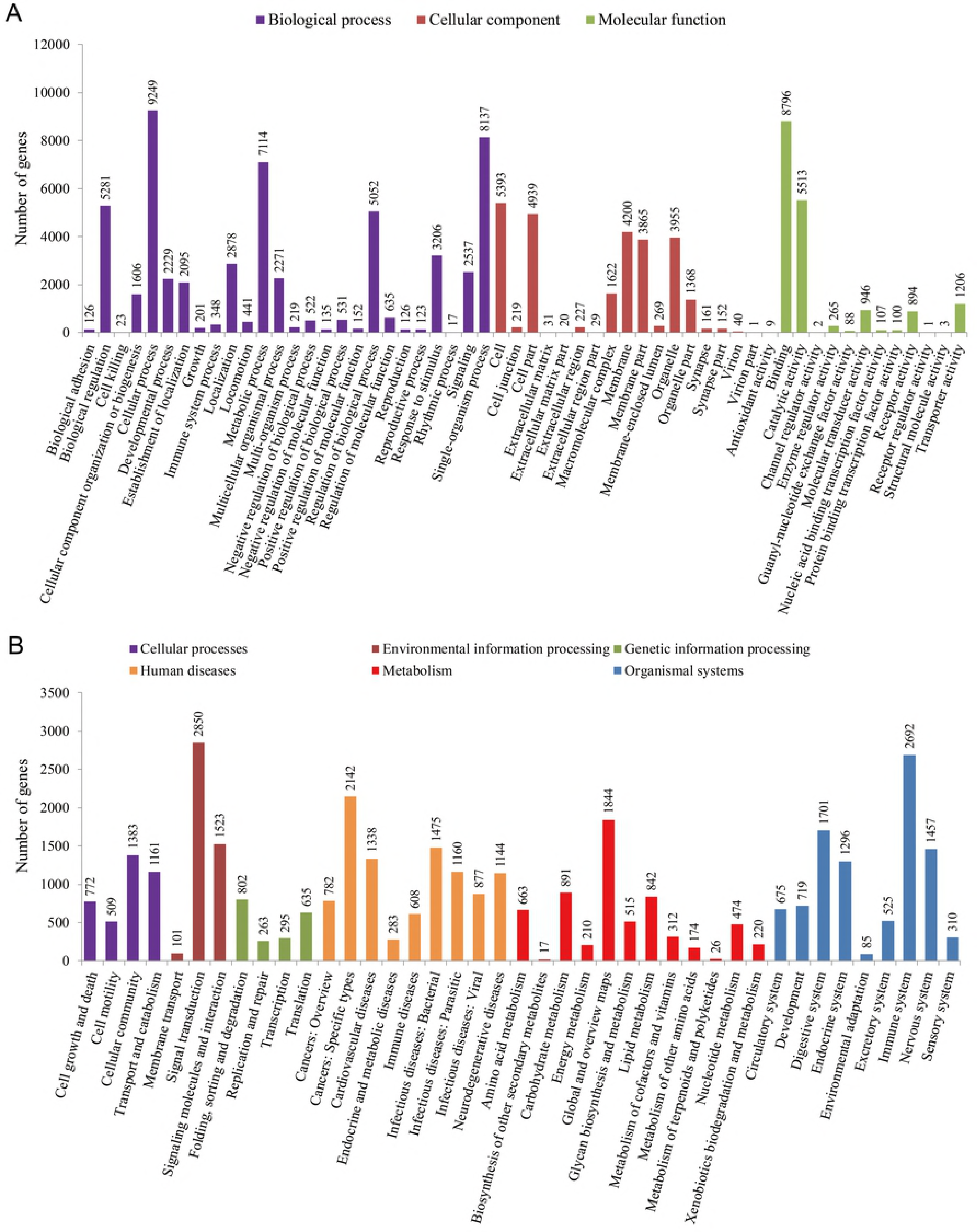
Gene Ontology (GO) and KEGG pathways functional classification of annotated unigenes. (A) A total of 16,954 unigenes showing significant similarity to homologous genes in GO databases were assorted to three main categories: cellular components, molecular functions, and biological processes. (B) KEEG pathway assignment based on six main categories: genetic information processing, metabolism, cellular processes, organismal systems, environmental information processing and human diseases.

Furthermore, all unigenes were also subjected to an analysis for possible functional prediction and classification by searching their predicted coding sequences of unigenes against KOG database. A total of 23,978 unigenes were successfully annotated and clustered into 25 subcategories (Fig 4, S4 Table). The largest orthology cluster with designation of ‘signal transduction mechanisms’ accounted for 24.42% (13,064) of the overall annotations, followed by ‘general function prediction only’ (8,968; 16.76%), ‘posttranslational modification, protein turnover, chaperones’ (4,280; 8.00%), ‘transcription’ (3,283; 6.14%), ‘intracellular trafficking, secretion, and vesicular transport’ (2,809; 5.25%),‘function unknown’ (2,695; 5.04%), ‘cytoskeleton’ (2,659; 4.97%), and ‘inorganic ion transport and metabolism’ (2,117; 3.96%). In summary, these functional annotation and classification would provide plentiful information resource for gene mining and function analysis.

**Fig 4.**
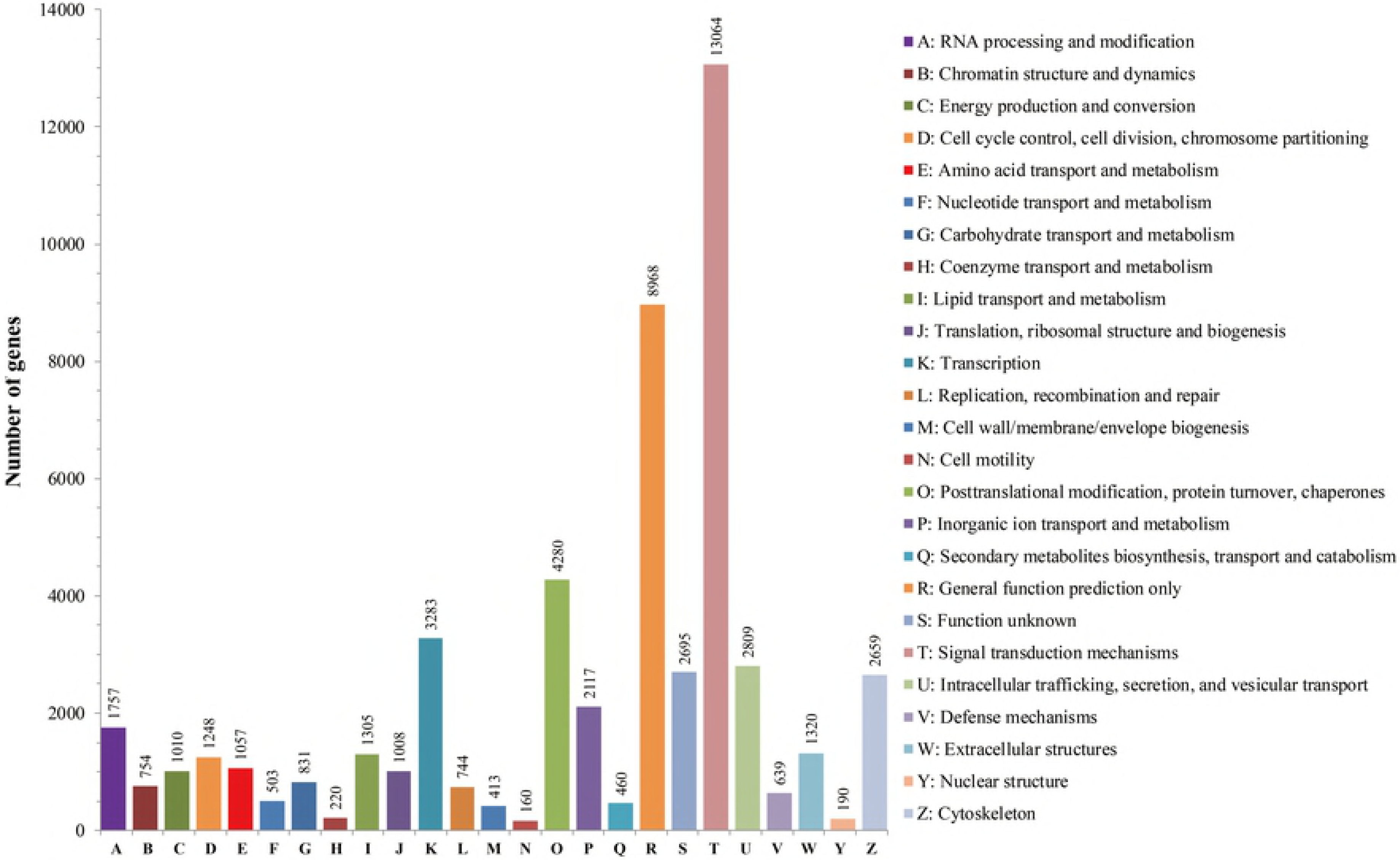
EuKaryotic orthologous group (KOG) classification of annotated unigenes. Approximately 23,978 of the 35,707 unigenes with Nr hits were grouped into 25 KOG categories.

### Differential expression analysis

In this study, SBGs were identified from the 69,836 unigenes by filtration according to criterion for differential expression. The analysis resulted in 17,683 (25.32%) unigenes differentially expressed between males and females (S5 Table, S1 Fig). These SBGs were further analyzed by their expression levels and the results suggested that most of them were up-regulated in males. Particularly, 11,875 SBGs showed higher expression in males (male up-regulated), while 5,808 SBGs showed higher expression in females (male down-regulated). Within the identified SBGs, 1,281 were specially expressed in males and 488 were specially expressed in females (S5 Table).

### Screening of SBGs potentially involved in aggressive behavior modulation

Based on the annotation and classification information, GO terms and KEGG pathways were further analyzed to screen out candidate SBGs associated with aggressive behavior modulation. The target pathways and terms mainly included ‘neuroactive ligand-receptor interaction’ (ko04080), ‘neurotrophin signaling pathway’ (ko04722), ‘estrogen signaling pathway’ (ko04915), ‘steroid hormone biosynthesis’ (ko00140), ‘receptor activity’ (GO:0004872), ‘reproduction’ (GO:0000003), ‘hormone activity’ (GO:0005179), ‘steroid hormone receptor activity’ (GO:0003707), ‘steroid biosynthetic process’ (GO:0006694) (Table 2).

**Table 2.** KEGG pathways and GO terms for identification of SBGs involved in aggressive behavior modulation.

| Database | Terms / Pathways | GO / KEEG ID | Number of transcripts | Total |
| --- | --- | --- | --- | --- |
| KEEG | Neuroactive ligand-receptor interaction | ko04080 | 629 | 1445 |
|  | Neurotrophin signaling pathway | ko04722 | 297 |  |
|  | Steroid hormone biosynthesis | ko00140 | 74 |  |
|  | Steroid biosynthesis | ko00100 | 26 |  |
|  | Endocrine and other factor-regulated calcium reabsorption | ko04961 | 158 |  |
|  | GnRH signaling pathway | ko04912 | 261 |  |
| GO | Receptor activity | GO: 0004872 | 915 | 1186 |
|  | Receptor regulator activity | GO: 0030545 | 1 |  |
|  | Reproduction | GO: 0000003 | 126 |  |
|  | Reproductive process | GO: 0022414 | 123 |  |
|  | steroid hormone receptor activity | GO:0003707 | 3 |  |
|  | hormone activity | GO:0005179 | 8 |  |
|  | steroid biosynthetic process | GO:0006694 | 10 |  |

Finally, a set of candidate SBGs with regulative roles for aggressive behavior were identified from the transcript data. Considering the functional differences, these SBGs were generally clustered into two categories: neurotransmitter and neuroendocrine pathways and sex steroid hormone pathways (Table 3). Most of SBGs showed higher expression levels in male *B. splendens*. Herein, the highly representative genes included male up-regulated SBGs such as 5-hydroxytryptamine receptors (*htr*), dopamine receptors (*drd*), γ-aminobutyric acid receptors (*gabr*), cholesterol side chain cleavage cytochrome P450 (*cyp11a1*), steroid 17-alpha-hydroxylase/17,20 lyase (*cyp17a1)*, testosterone 17-beta-dehydrogenase 3 (*hsd17b3*), doublesex and mab-3 related transcription factor 1 (*dmrt1*), nuclear receptor subfamily 0 group B member 1 (*dax1*), steroidogenic factor 1 (*sf-1*), and female up-regulated SBGs such as 3-keto-steroid reductase (*hsd17b7*), gonadal soma derived factor 1 (*gsdf1*) and protein fem-1 homolog C (*fem1c*).

**Table 3.** SBGs putatively involved in aggressive behavior modulation in *B. splendens*.

| Gene name | Nr annotation | E-value | Transcript ID | Length (bp) | Log<sub>2</sub>Fold Change (♂/♀) | FDR | Trend |
| --- | --- | --- | --- | --- | --- | --- | --- |
| **Neurotransmitter and neuroendocrine pathways** |  |  |  |  |  |  |  |
| *htr2a* | 5-hydroxytryptamine receptor 2A-like | 0 | Unigene0030158 | 4450 | 1.961 | 1.10E-149 | UP |
| *htr3a* | 5-hydroxytryptamine receptor 3A-like | 3E-164 | Unigene0031141 | 2053 | 5.714 | 1.04E-107 | UP |
| *htr1e* | 5-hydroxytryptamine receptor 1E | 0 | Unigene0039721 | 1422 | 3.089 | 4.78E-39 | UP |
| *htr3c* | 5-hydroxytryptamine receptor 3C-like | 2E-138 | Unigene0068687 | 986 | 1.789 | 3.33E-13 | UP |
| *htr4* | 5-hydroxytryptamine receptor 4 | 2E-77 | Unigene0052106 | 382 | 4.456 | 2.93E-10 | UP |
| *SLC6A4* | Sodium-dependent serotonin transporter-like | 7E-84 | Unigene0041170 | 413 | 2.496 | 0.000339899 | UP |
| *drd5* | D(1B) dopamine receptor-like | 8E-108 | Unigene0046850 | 667 | 1.385 | 0.000952899 | UP |
| *drd2a* | D(2) dopamine receptor A-like | 3E-162 | Unigene0031216 | 854 | 1.478 | 6.67E-11 | UP |
| *drd2* | D(2)-like dopamine receptor-like | 1E-80 | Unigene0009674 | 917 | 3.073 | 6.19E-26 | UP |
| *hrh1* | Histamine H1 receptor | 3E-74 | Unigene0042840 | 403 | 1.648 | 0.02659 | UP |
| *hrh2* | Histamine H2 receptor-like | 6E-166 | Unigene0045571 | 2010 | -6.249 | 0 | Down |
| *hrh3* | Histamine H3 receptor-like | 8E-18 | Unigene0027814 | 490 | 2.192 | 8.29E-05 | UP |
| *hrh4* | Histamine H4 receptor | 5E-39 | Unigene0006615 | 251 | 2.522 | 0.044731 | UP |
| *gabra4* | Gamma-aminobutyric acid receptor subunit alpha-4 | 2E-32 | Unigene0038838 | 543 | 2.233 | 0.000910315 | UP |
| *gabra5* | Gamma-aminobutyric acid receptor subunit alpha-5-like | 0 | Unigene0043252 | 5187 | 1.034 | 1.39E-94 | UP |
| *gabrr3* | Gamma-aminobutyric acid receptor subunit rho-3-like | 1E-09 | Unigene0003367 | 454 | 1.510 | 0.00695924 | UP |
| *gabbr1* | Gamma-aminobutyric acid type B receptor subunit 1-like | 0.000004 | Unigene0052902 | 325 | 4.920 | 2.36E-07 | UP |
| *gabbr2* | Gamma-aminobutyric acid type B receptor subunit 2-like | 1E-26 | Unigene0030612 | 269 | 2.969 | 0.00571534 | UP |
| *mtnr1b* | Melatonin receptor 1B | 3E-110 | Unigene0068593 | 511 | 1.726 | 0.00057695 | UP |
| *avpr* | Arginine vasotocin receptor | 5E-138 | Unigene0001414 | 637 | 1.098 | 0.0103464 | UP |

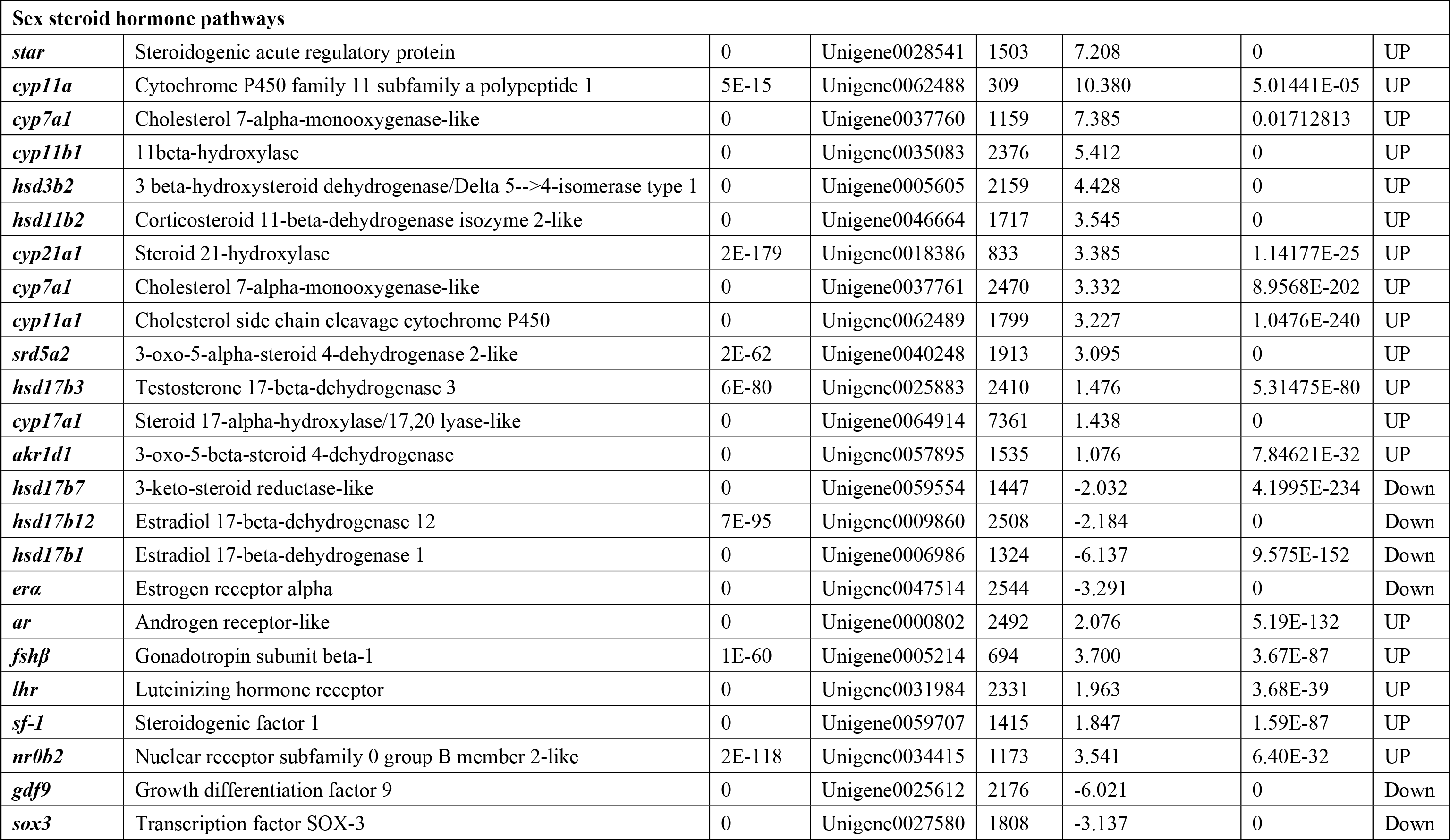

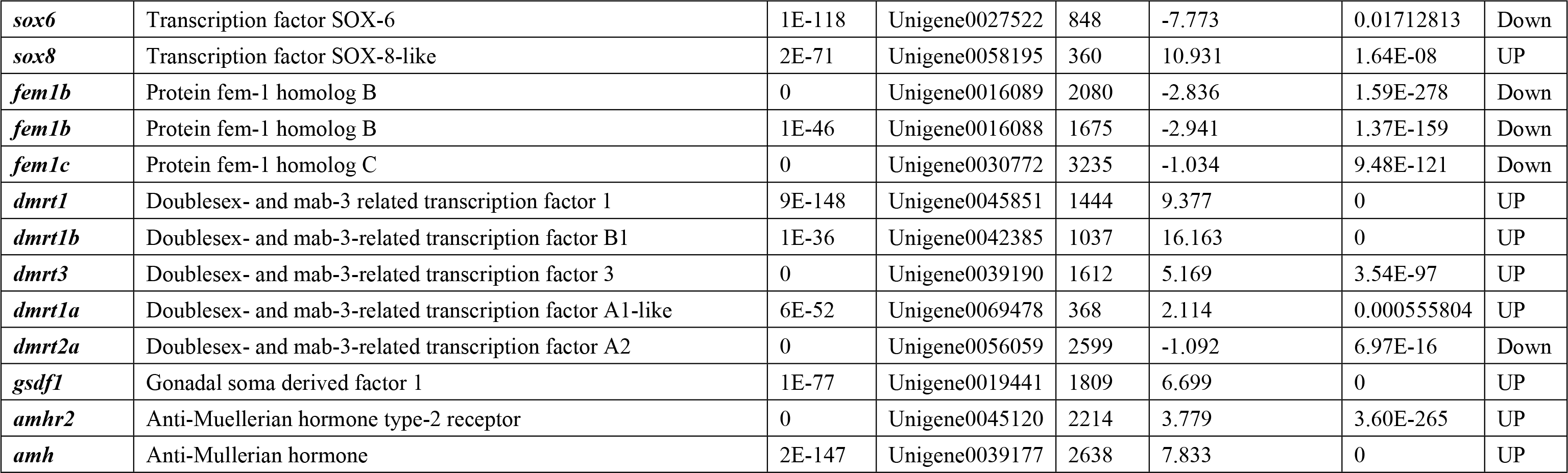

### Validation of expression level of SBGs by qRT-PCR

Twelve SBGs of interest were chosen and validated by qRT-PCR, including seven male-biased genes (i.e. *htr3a*, *hsd3b2*, *cyp11a1*, *cyp17a1*, *ar*, *gsdf1*, *amh*) and five female biased genes (i.e. *hrh2*, *erα*, *gdf9*, *hsd17b7*, *fem1c*). The validation results demonstrated that almost all genes displayed similar expression patterns both in RNA-Seq and qRT-PCR (Table 4). Meanwhile, a correlation analysis was conducted between the RNA-Seq data and qRT-PCR expression data (Fig 5). The consistent tendencies of expression levels between RNA-Seq and qRT-PCR data (*R*^2^ = 0.9088) confirmed the accuracy of gene expression levels quantified by RNA-Seq.

**Table 4.**
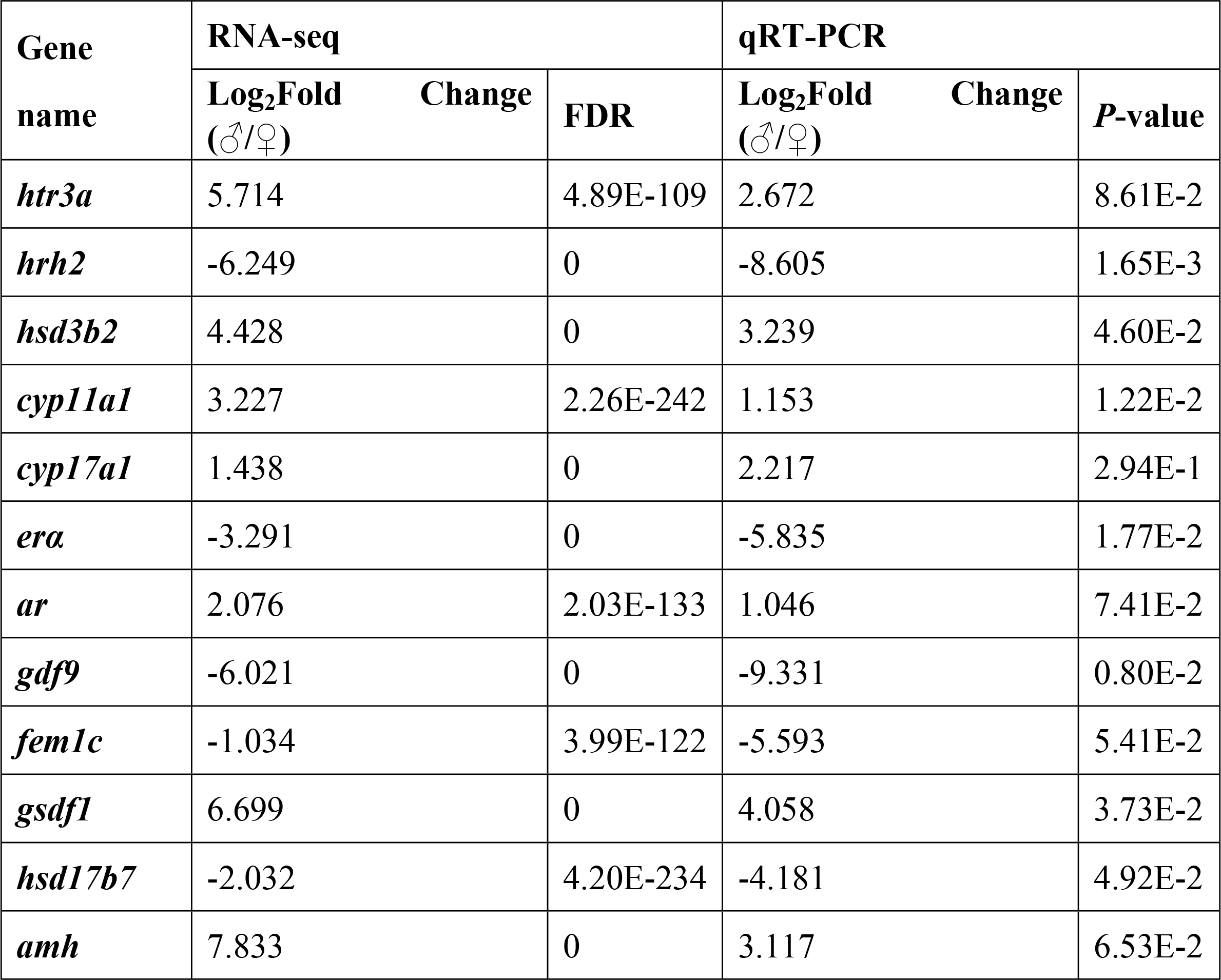
Validation of the RNA-seq data by qRT-PCR analysis.

**Fig 5.**
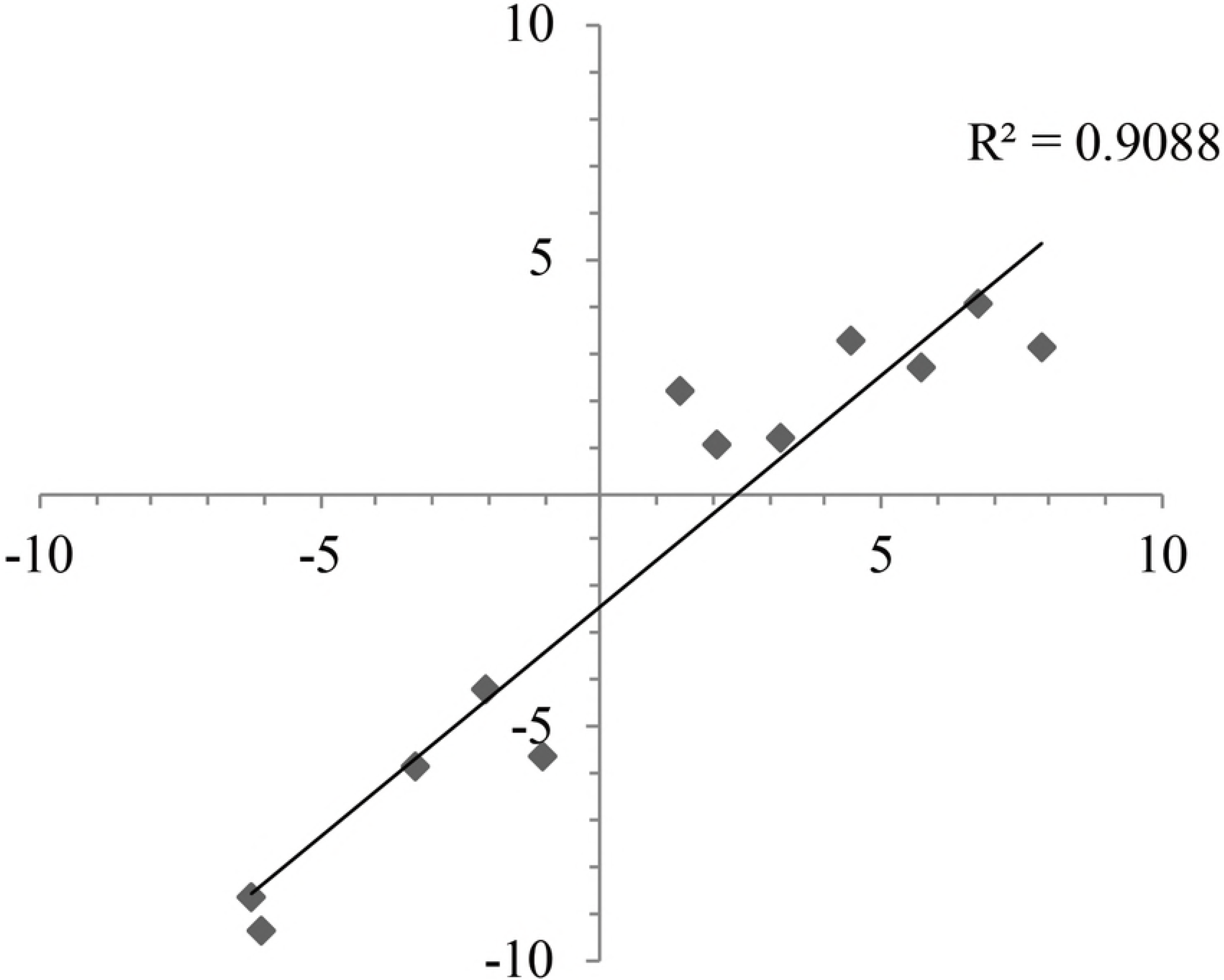
Differential expression validation of SBGs by qRT-PCR. The consistency of log_2_(Fold Change) between RNA-Seq data (X-axis) and qRT-PCR analysis (Y-axis) is high (R^2^= 0.9088) based on the selected SBGs.

### Characterization of SSR loci and polymorphism validation

Using the MISA software, a total of 9,617 (13.77%) unigene sequences containing 12,751 potential SSRs were detected, with 1,809 sequences contained more than one SSR locus (Fig 6A). The SSR distribution density in transcriptome was one SSR loci per 5.99 kb. The most abundant SSRs were di-nucleotide repeats (6,418; 50.33%), followed by tri-nucleotide repeats (4,657; 36.52%), tetra-nucleotide repeats (1,209; 9.48%), penta-nucleotide repeats (279; 2.19%) and hexa-nucleotide repeats (188; 1.47%) (Fig 6B). The copy number of different repeat units ranged from 4 to 41. A further analysis on the frequency distribution of motif sequence types was conducted. Within the di- and tri-nucleotide repeats motifs, AC/GT (4,445; 34.86%), AG/CT (1,466; 11.50%) and AGG/CCT (1,297; 10.17%) repeats were the three predominant types in the *B. splendens* transcriptome (Fig 6C).

**Fig 6.**
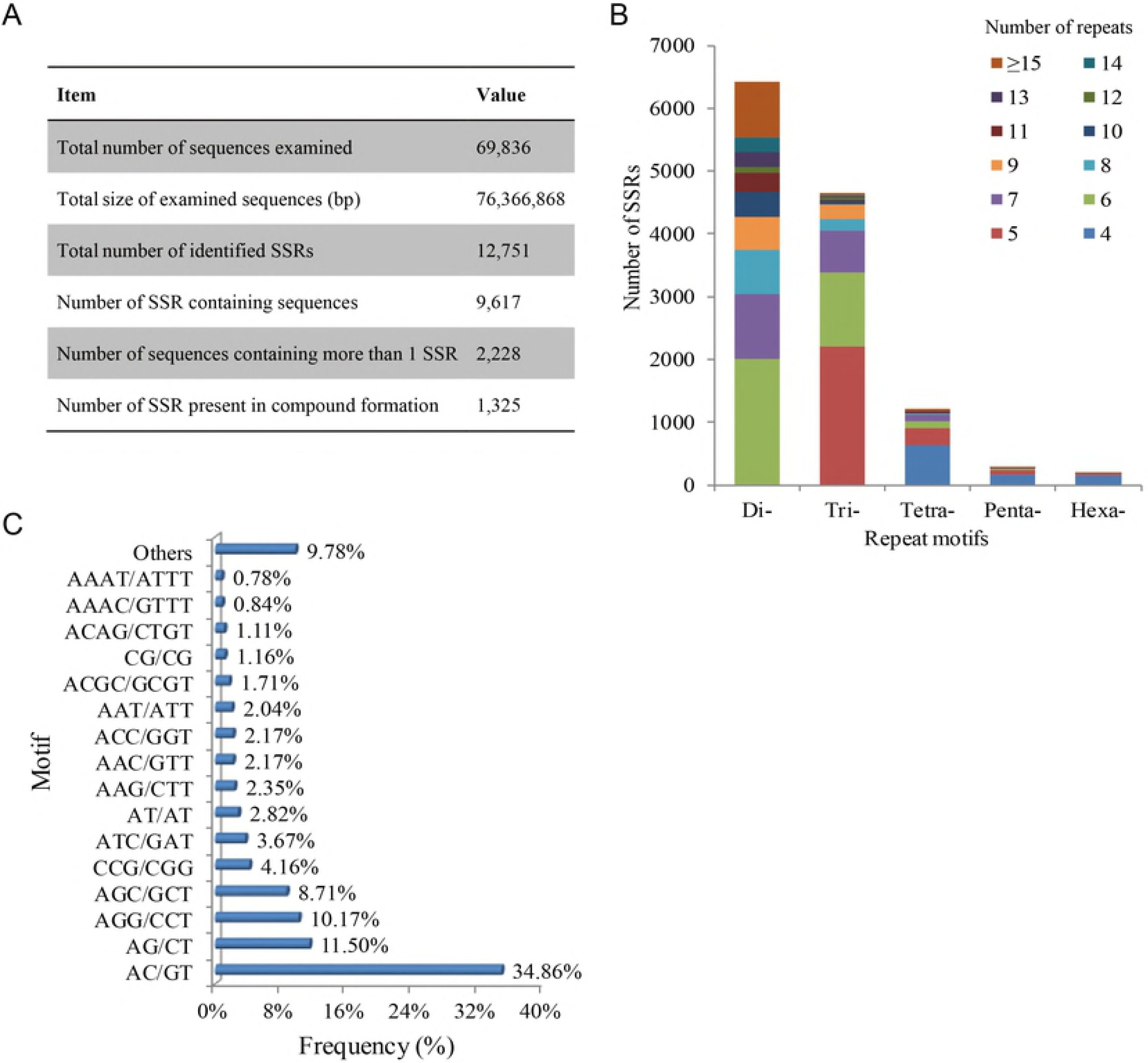
Characterization of SSR loci identified in the *B. splendens* transcriptome. (A) Statistics of SSR searching results. (B) The distribution of repeat number for di-, tri-, tetra-, penta-, and hexa-nucleotide SSR repeat motifs. (C) The frequency distribution of classified types of repeat motifs; The frequency of main motif types was displayed.

For microsatellite marker development, 7,970 (62.50%) SSR-containing sequences enabled the design of primers. The information of all SSR primer pairs derived from unigenes were listed in S6 Table. In order to assess the genetic polymorphism of these markers, one hundred pairs of SSR primers were randomly selected for primer synthesis and PCR validation. As a result, 53 primer pairs successfully generated stable and reproducible target amplification products using genomic DNA. For these 53 primer pairs, a *B. splendens* population containing 30 individuals was analyzed. Thirty-four SSR loci were monomorphic and 19 loci were polymorphic. Of these polymorphic loci, a total of 65 alleles were successfully amplified and the number of alleles per locus ranged from 2 to 5, with an average of 3.42. The *H*_o_ and *H*_e_ values ranged from 0.167 to 0.933 and from 0.430 to 0.772, with an average of 0.575 and 0.618, respectively. The *PIC* values varied from 0.357 to 0.617, with a mean value of 0.534 (Table 5).

**Table 5.**
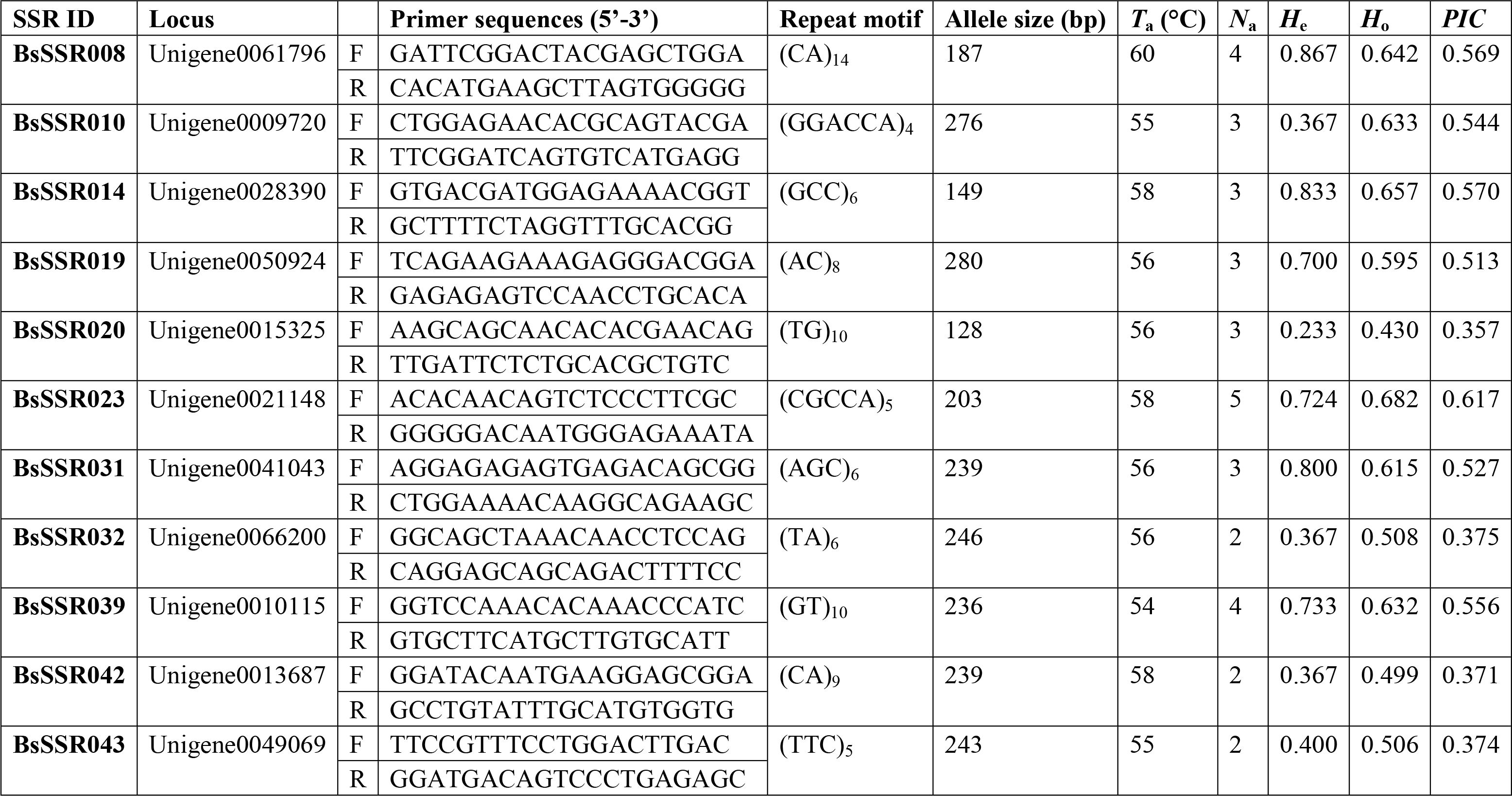
Characterization of 19 polymorphic SSR loci in 30 *B. splendens* individuals.

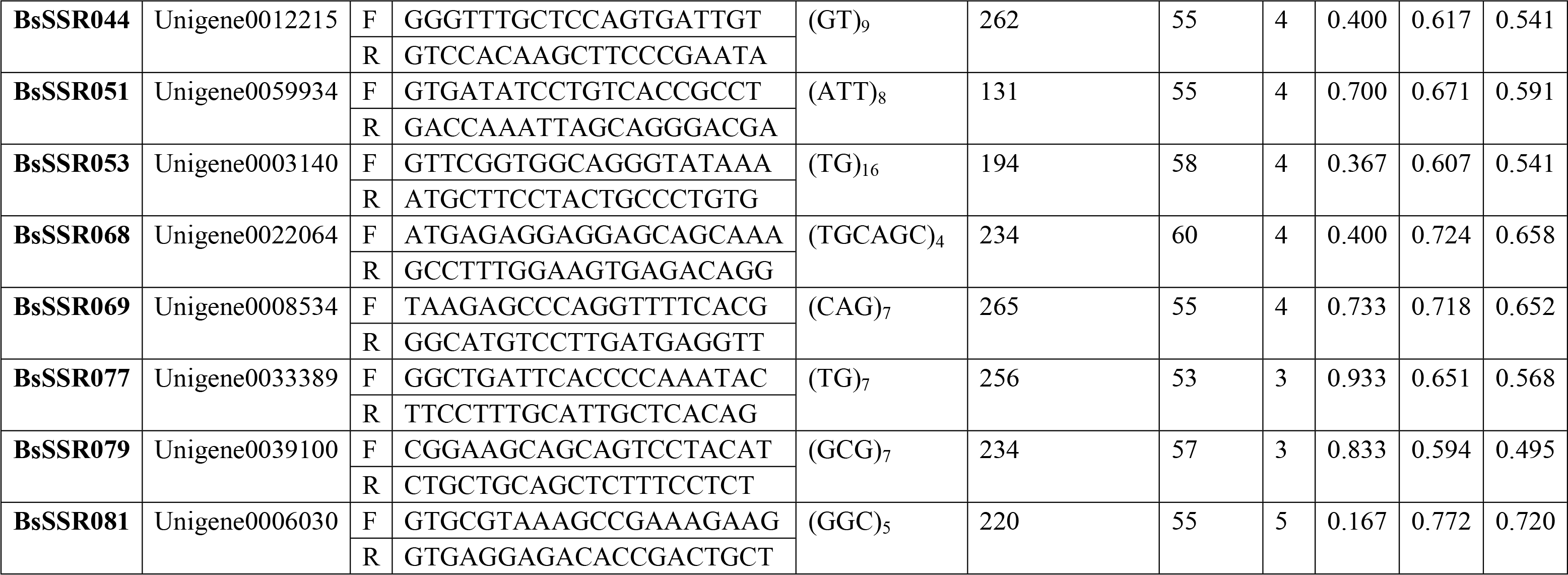

## Discussion

### Characterization of the *B. splendens* transcriptome

Transcriptome sequencing is an increasingly preferred choice to obtain large-scale functional gene sequences and to enrich the genetic resource database rapidly and cost-effectively for non-model species [36]. In order to generate a representative transcriptome of *B. splendens*, different tissues were sampled for RNA isolation. The extracted total RNAs were pooled in equal quantities for male and female cDNA libraries construction, respectively. Such multi-tissues strategy has been widely employed in the RNA-Seq projects for other teleosts [27, 37–39]. For high-throughput short read sequencing, assembly with high-quality will provide benefit for the post transcriptomic analysis like annotation, gene identification and comparative genomics. According to most of published *de novo* transcriptome assembly, the quality of assembly is principally evaluated by the length distribution of transcripts [40]. In the present study, more than half of the *de novo* assembled unigenes were greater than 500 bp in length and the mean length of unigenes reached 1093 bp (Table 1, Fig 1). Several similar results were also given by Trinity transcriptome assemblies for teleost fishes such as *Pelteobagrus fulvidraco* (1611 bp) [41], *Trachinotus ovatus* (1179 bp) [37] and *Scatophagus argus* (906 bp) [27], strongly indicating that our transcriptome data was assembled effectively. In contrast with other *de novo* assembly softwares e.g. Newbler [42], iAssembler [43] and CLC Genomics Workbench [44], the most satisfying assembler Trinity could provide a unified solution for good-quality transcriptome reconstruction in species without a reference genome [26, 40].

A longer assembled sequence can provide more adequate information for further gene investigation and facilitate identification of molecular markers. In this research the BLAST match rates of unigenes enhanced markedly from 200 - 300 bp (32.82%) to 1100 - 1200 bp (80.74%) in length. On the other hand, the query sequence length was crucial for determining the level of significance of the BLAST hits. The ratio of unigenes with significant BLAST scores increased sharply from 200 - 500 bp to 500 - 1500 bp. Taken together, these results indicate that the proportion of sequences with matches in database is greater among the longer assembled sequences, which is in agreement with other analytical outcomes of next-generation transcriptome sequencing [27, 45, 46]. For functions prediction of the transcript sequences, 51.19 % of the unigenes were matched to homologous sequences by searching against public databases (Fig 2A), suggesting the successful annotation of the transcriptomic sequences. However, we notice that approximate half of transcripts (48.81%) failed to be annotated to any sequences in the queried databases. Ground on previous studies, a high percentage of unannotated sequence is generally caused by the un-translated mRNA regions, chimeric sequences with assembly errors, non-conserved protein regions or novel genes [39]. It is unsurprising that the low annotation ratio seems to commonly occur in non-model animals without published genomes, especially in aquatic species [37, 39, 47].

### SBGs involved in aggression behavior modulation

Previous studies have implicated neurotransmitter pathways as critical roles in controlling aggression behavioral syndrome in mammals, birds and fishes [48]. Serotonin (5-HT), dopamine, γ-aminobutyric acid (GABA) and histamine systems are generally thought to represent the key components of neuroregulatory network in aggressive display. Of which, 5-HT plays a more important role in aggressive behavior modulation through binding to a diverse class of receptors in a wide range of species [49]. Extensive evidence has shown that 5-HT exerts a primarily inhibitory effect on aggression expression by receptor activation [50, 51], including treatment with 5-HT reuptake inhibitors [8] or receptor selective agonists [10, 49]. Actually, neural pathways can mediate different reactions depending on the involved receptor subtypes. The inhibitory nature of 5-HT system has been attributed to stimulation of the 5-HT type 1 (5-HT_1A_, i.e. 5-HT_1A_ and 5-HT_1B_) [10, 52, 53] and 5-HT_2_ receptors [54, 55]. 5-HT could decrease aggressive behavior via 5-HT_1A_ receptors in *B. splendens* [10]. By contrast, activation of the 5-HT_3_ receptor has been linked to facilitating aggression [56, 57]. It has been proved that 5-HT_1B_ receptor can actually induce aggressive behavior instead of inhibiting it [58]. All above results suggest a complex role for 5-HT system in the expression of aggression. Despite the large amount of research on the linkage between 5-HT and aggression, their intricate relationship leads to the unclearness of the precise roles. In our current study, we identified five male-biased 5-HT receptor subtype genes (i.e. *htr2a*, *htr3a*, *htr1e*, *htr3c*, *htr4*) by transcriptome-based gene mining (Table 3). Currently none of these receptors has been subjected to any investigation on aggressive behavior modulation in teleosts. Hence the candidate transcripts would make good starting points for further research into the mechanisms underlaying the aggression control of 5-HT system.

A quantity of SBGs encoding dopamine, GABA and histamine receptors were obtained by differential expression analysis, such as *drd2*, *drd5*, *hrh1*, *hrh2*, *gabbr1* and *gabbr2* (Table 3). The dopaminergic system generally plays an active role in the modulation of aggression. Hyperactivity in the dopamine system is substantially linked with increases in aggressive behaviors [59, 60]. Among the five dopamine receptor families, dopaminergic manipulations with D_1_, D_2_ and D_3_ receptor antagonists can decrease aggressive behavior [61, 62]. Nevertheless, dopamine can sometimes reduce the impulsivity that might lead to abnormal aggression [63]. As a main inhibitory neurotransmitter, the relationship of GABA with aggression is extremely complex and highly associated with 5-HT. GABA_a_ receptor activation is associated with a decrease in aggressive behavior [63, 64]. By contrast, GABA_b_ receptors are directly related to escalated aggressive behavior [65]. Moreover, histamine system has been demonstrated to take part in the aggressive display. Certain changes in aggressive behavior have been observed following histamine administration and distinct roles for H_1_ and H_2_ receptors have been delineated [66]. Reduced brain histamine levels increase aggression-boldness in adult zebrafish, and this behavioral phenotype can be rescued by pharmacological increase of histamine signaling [48]. It is very interesting that almost all of these receptor SBGs, except *hrh2*, were up-regulated in male *B. splendens* (Table 3), suggesting that such neurotransmitter systems may play certain roles in the specific expression of male aggression. But their exact effects involved in aggressiveness are still far from being fully elucidated, especially in teleosts. To improve our knowledge on the neurobiological bases of aggressive behavior, future research should also be carried out to investigate the nature of interactions among these neurotransmitter systems.

Sex steroid hormones are generally believed to play key roles in behavioral control. In our study, a number of SBGs participating in sex steroid hormone pathways were identified. They are comprised of steroidogenic genes, sex hormone receptor genes and regulatory factors for steroidogenensis (Table 3). The classic association between sex steroid hormones (i.e. androgens, estrogens) and aggression has been confirmed in many vertebrate species [67–71]. Androgens such as testosterone and 11-ketotestosterone have been demonstrated to mediate the expression of intra-sexual aggression in teleost [72]. In testosterone manipulated sex reversal in female *B. splendens*, aggression increased toward males as treatment progressed [73]. In contrast, many reported research has revealed that 17β-estradiol (E_2_) or estrogen mimics (e.g. 17α-ethinylestradiol [23, 74], phytoestrogens [21], benzophenone-3 [75]) can inhibit aggression of males by disrupting steroidogenesis of sex hormones. *B. splendens* exposed to E_2_ showed reduction in aggressive behavior and displayed higher frequency of inactive behaviors [76]. It is well known that steroid hormones regulate aggressive behaviors by binding to receptors in the central nervous system. Some findings in rodents indicate that the neural androgen receptor (AR) is required for mediating activational effects of testosterone in the regulation of aggressive behavior [77, 78]. Previous studies revealed that the lack of estrogen receptor (ER)-α gene severely reduced the induction of male aggression, while mice that lacked the ER-β gene tended to be more aggressive [68, 78, 79]. In this study most of the steroidogenic SBGs (e.g. *cyp17a1*, *cyp11a1* and *hsd3b2*) are male up-regulated genes that participate in the androgen biosynthesis (Table 3). These findings implying that androgen may have a central role in the regulation of male aggression expression in *B. splendens*. To our knowledge, however, no study has been done to address the actions of these genes and little is known about their effects in aggressive regulation. With the aid of the available transcript resources, more well-designed molecular biology and biochemical studies focused on steroid hormone pathways are absolutely essential and highly encouraged to unravel the in-depth regulatory mechanism underlying aggression expression.

### EST-SSR characterization and marker validation

In the *B. splendens* transcriptome, about 13.77% of assembled unigenes were identified as SSR-containing sequences (Fig 6A). The proportion is similar to that in *S. argus* [27], higher than that in *Megalobrama amblycephala* (5.0%) [80], but considerably lower than that in *Paralichthys olivaceus* (22.14%) [81], *Sander lucioperca* (29.0%) [82] and *Pelteobagrus fulvidraco* (49.0%) [83]. Multiple reasons that are likely able to explain this variance include the nature of different species and potential artifacts from the usage of different software tools. Within the SSR-containing unigenes, 2,228 contained more than one SSR and 255 contained more than two SSRs. The inequality of distribution of SSRs has been also detected in transcriptome-based SSR development for other fishes such as *S. argus* [27], *Acipenser sinensis* [84] and *Carassius auratus* [85]. On average, one SSR could be found every 5.99 kb in *B. splendens* transcriptome. The distribution density of microsatellites was similar to reports for other teleosts such as *T. ovatus* [37] and *Salmo trutta* m. *trutta* [44]. Additionally, di-nucleotide (AC/GT, AG/CT) and tri-nucleotide repeats (AGG/CCT) were found to be the most abundant SSR motifs (Fig 6 B,C). Similar results were observed in the transcriptomic analysis of diverse species of fish including *T. ovatus* [37], *Scophthalmus maximus* [39], *S. argus* [27] and *S. trutta* m. *trutta* [44]. Overall, such consistence implicates the conservativeness of microsatellite distribution and dominating repeat type of SSR motif in teleosts.

Amount to 9,617 unigenes possessing SSR loci were identified and 7,970 EST-SSR markers were developed (Fig 6A). In order to evaluate the quality of the EST-SSR primers designed in this study, we analyzed 100 randomly selected primers and found that 53 primer pairs could successfully amplified their target amplicons (53% success rate). This result means that approximately one-half of the 7,970 SSR primers are expected to amplify successfully their targets. Here about half of the primer pairs might have been mismatched and, in such cases, some EST-SSRs may have failed to amplify because large introns were present in the target amplicon, or because the designed primers were across splice sites. Nearly 20% of the primer pairs specifically yielded target PCR products and exhibited polymorphisms (Table 5), demonstrating that we have successfully achieved the aim of development of large scale SSR markers for *B. splendens*. The rate of polymorphic EST-SSRs isolated in this study was slightly lower than that in some other fishes [39, 86, 87], perhaps because the tested individuals came from the same fighting fish hatchery center. More polymorphic microsatellites would be developed if more geographically distant populations were subjected to the polymorphism validation.

As tandemly repeated motifs with high-levels of polymorphism, SSRs are widely used as molecular markers in various aspects of molecular genetic studies. Owing to the availability of genomic and transcriptome sequences base on NGS platforms, *de novo* screening for large sets of SSRs has become more economic and efficient in non-model organisms. Transcriptome- and EST-based SSRs mainly occur in the coding regions of annotated genes, they are important resource for determining functional genetic variation [44]. EST-SSR marker has become an effective approach to identify phenotypic associations in fish species. They are useful resources for genetic map construction, map-based gene clone, comparative genomics, genetic diversity analysis, functional genetic variation examination and molecular marker assisted selection [88]. To date, only a few SSR markers have been developed for *B. splendens*. The potential EST-SSRs obtained here would be a wealth of resource for developing polymorphic markers, which could be utilized as a powerful tool for future studies on genetics and genomics.

## Conclusions

In this study, we used Illumina high-throughput sequencing to analyze the comprehensive transcriptome of *Betta splendens*. An informative transcriptomic dataset composed of 69,836 transcripts was achieved. This transcriptome greatly increases the available genetic information for general gene expression studies and contributes valuable sequence database for functional gene discovery and analysis. Here it especially facilitates the characterization of sexual dimorphism of regulatory gene in aggressive behavior expression. A number of candidate sex-biased genes involved in aggressive modulation will provide good starting points for future studies on the regulatory mechanisms underlying aggression display. Finally, a large amount of SSRs was detected, supplying abundant marker resources for future research into molecular genetics and genomic studies.

## Acknowledgments

We thank Mr. Chenggui Wang and Ms. Mei Wang (Guangdong Ocean University, China) for help with experiments. We also thank Ms. Haiyu Jing and Ms. Liu Yang (Gene Denovo Biotechnology Co., Ltd, China) for technical support in Illumina sequencing and bioinformatic analysis.

## Supporting Information

**S1 Table. The primers for qRT-PCR validation.**

(DOCX)

**S2 Table. The GO annotation results of the unigenes in the *B. splendens* transcriptome.**

(XLSX)

**S3 Table. KEEG pathway analysis for the transcriptome of *B. splendens.***

(XLSX)

**S4 Table. The functional classification of the KOG classes for the transcriptome of *B. splendens.***

(XLSX)

**S5 Table. The functional annotation of the differentially expressed unigenes between males and females.**

(XLSX)

**S6 Table. Information of all SSRs primer pairs derived from unigenes in the *B. splendens* transcriptome.**

(XLSX)

**S1 Fig. Differential expression analysis of unigenes between male and female *B. splendens.***

A total of 17,683 unigenes differentially expressed between males and females. Of which, 11,875 unigenes showed higher expression in males (red), while 5,808 unigenes showed higher expression in females (green). (TIFF)

**S2 Fig. PCR amplification profiles of 30 *B. splendens* accessions using primer pair BsSSR068.**

The PCR amplified products were separated on 8.0% polyacrylamide gel. M indicated the pBR322 DNA/MspI marker. (TIFF)

